# Carbohydrate complexity structures stable diversity in gut-derived microbial consortia under high dilution pressure

**DOI:** 10.1101/2020.01.31.929760

**Authors:** Tianming Yao, Ming-Hsu Chen, Stephen R. Lindemann

**Affiliations:** Whistler Center for Carbohydrate Research, Department of Food Science, Purdue University. 745 Agriculture Mall Drive, West Lafayette, IN 47907, USA; Department of Nutrition Science, Purdue University. 700 W. State Street, West Lafayette, IN 47907, USA

**Keywords:** dietary fiber, inulin, arabinoxylan, human gut microbiota, consortia

## Abstract

Dietary fibers are major substrates for the colonic microbiota, but the structural specificity of these fibers for the diversity, structure, and function of gut microbial communities are poorly understood. Here, we employed an *in vitro* sequential batch fecal culture approach to determine: 1) whether the chemical complexity of a carbohydrate structure influences its ability to maintain microbial diversity in the face of high dilution pressure and 2) whether substrate structuring or obligate microbe-microbe metabolic interactions (e.g. exchange of amino acids or vitamins) exert more influence on maintained diversity. Sorghum arabinoxylan (SAX, complex polysaccharide), inulin (low-complexity oligosaccharide) and their corresponding monosaccharide controls were selected as model carbohydrates. Our results demonstrate that complex carbohydrates stably sustain diverse microbial consortia. Further, very similar final consortia were enriched on SAX from the same individual’s fecal microbiota across a one-month interval, suggesting that polysaccharide structure is more influential than stochastic alterations in microbiome composition in governing the outcomes of sequential batch cultivation experiments. SAX-consuming consortia were anchored by *Bacteroides ovatus* and retained diverse consortia of >12 OTUs; whereas final inulin-consuming consortia were dominated either by *Klebsiella pneumoniae* or *Bifidobacterium* sp. and *Escherichia coli*. Furthermore, auxotrophic interactions were less influential in structuring microbial consortia consuming SAX than the less-complex inulin. These data suggest that carbohydrate structural complexity affords independent niches that structure fermenting microbial consortia, whereas other metabolic interactions govern the composition of communities fermenting simpler carbohydrates.

**IMPORTANCE:** The mechanisms by which gut microorganisms compete for and cooperate on human-indigestible carbohydrates of varying structural complexity remain unclear. Gaps in this understanding make it challenging to predict the effect of a particular dietary fiber’s structure on the diversity, composition, or function of gut microbiomes, especially with inter-individual variability in diets and microbiomes. Here, we demonstrate that carbohydrate structure governs the diversity of gut microbiota under high dilution pressure, suggesting that such structures may support microbial diversity *in vivo*. Further, we also demonstrate that carbohydrate polymers are not equivalent in the strength by which they influence community structure and function, and that metabolic interactions among members arising due to auxotrophy exert significant influence on the outcomes of these competitions for simpler polymers. Collectively, these data suggest that large, complex dietary fiber polysaccharides structure the human gut ecosystem in ways that smaller and simpler ones may not.

## INTRODUCTION

Gut microbiota play an increasingly appreciated role in the human digestive system and are now regarded as an important “forgotten organ” in maintaining health and treating disease (1). For instance, the gut microbiota convert dietary fibers into various bacterial metabolites, including short-chain fatty acids (SCFAs), that influence diverse health properties (2). Altered microbiome structures are associated with chronic diseases such metabolic syndrome, diabetes, and inflammatory bowel diseases (3–5); disturbed intestinal ecosystems (6–8) and chronic disease (9) are both associated with low diversity. However, the mechanisms that maintain diversity in the human gut microbiome are poorly understood but of great potential interest to modulate health.

Environmental factors, such as eating and fasting cycles, lifestyle shifts, infections, age and diet, can alter the composition of gut microbiota. Dietary fibers, which are carbohydrates resistant to human digestive enzymes and which escape small intestinal degradation and enter the colon, strongly influence the composition and function of gut microbiota (10). The capability of a gut microorganism to consume a dietary fiber depends on 1) its carbohydrate-active enzyme (CAZyme)-encoding genes that are specific to cleavage of different carbohydrate linkages (11), 2) its genome-encoded suite of transporters that take up the products of polysaccharide hydrolysis for use as carbon and energy sources, and 3) how those genes are regulated (12).

The competitive exclusion principle of ecology (Gause’s Law) states that two species competing for the same limiting resource cannot coexist under stable environmental conditions (13, 14). In the colonic environment, microbiota compete for non-digestible carbohydrates, but the effect of carbohydrate structure on ecological outcomes of competition are poorly understood. As CAZymes are highly specific for their cognate substrates and carbohydrates are very structurally diverse (15), multiple CAZymes are required to hydrolyze complex polysaccharides. Consequently, it is possible that organisms may specialize in consuming specific carbohydrate linkages, dividing metabolic labor and promoting coexistence through niche differentiation. However, many organisms (e.g., *Bacteroides* sp.) have the genetic potential to hydrolyze all glycosidic bonds in some carbohydrates (16, 17), suggesting the possibility that such generalists may successfully exclude others in competition for these resources. In this study, we hypothesized that carbohydrate structural complexity would maintain diverse gut microbiota. Specifically, we hypothesized that the more-complex the carbohydrate structure, in terms of the number of enzymes required for a polymer’s degradation, 1) the more likely the polymer will impart strong selective pressure on community structure and, 2) the greater the sustained diversity of species under high dilution pressure. To test this hypothesis, we selected inulin as a model simple carbohydrate (both small in size and requiring only a single hydrolase for degradation) and sorghum arabinoxylan (SAX) as a model complex polysaccharide (a high molecular-weight polymer with diverse glycosyl residues and linkages among them).

To identify the microbial populations most efficient for consumption of each substrate, we employed sequential batch fermentation; consortia were continuously passaged under high dilution pressure (1:100 dilution per day) in media containing model carbohydrate substrates differing in complexity. These were then compared to identical cultures fermenting monosaccharide controls mimicking the sugar composition of the respective polymers. Sequential batch fermentation strongly selects for the organisms with the highest growth rates under experimental conditions, rapidly diluting out slow- or non-growing organisms. Further, we investigated whether carbohydrate structure or other interactions were more influential in structuring the resulting communities via the addition of exogenous nutrients to alleviate potentially obligate metabolic interactions arising from auxotrophy. Our results support the hypothesis that increasing carbohydrate complexity affords greater numbers of independent niches, which select for stable, carbohydrate-specific consortia. Furthermore, our data suggest that increasingly complex carbohydrates (i.e. SAX) more strongly govern the composition of fermenting microbial consortia than simpler ones (i.e. inulin), which are more strongly governed by other metabolic interactions. This study reveals that complex carbohydrates effectively structure gut microbial consortia and maintain diversity against strong dilution pressure.

## RESULTS

### Carbohydrate polymer structural properties

The structural complexity of inulin and SAX with respect to glycosyl residue composition and molecular sizes are shown in Table 1 and Supplementary Figure 1. Fructose was the major component in the inulin employed in this study (86.2%) with a small amount of terminal glucose (13.8%). In contrast, SAX was predominantly composed of arabinose (35.1%) and xylose (37.0%), with minor components of glucose (7.6%), galactose (6.0%), and mannose (4.0%). Arabinoxylans (AXs) are hemicelluloses and are typically composed of β-1,4-linked xylose backbones with substituted arabinose branches. The arabinose/xylose ratio was 0.95, indicating SAX was highly branched. In size exclusion chromatography (Supplementary Figure 1), SAX eluted much faster than the inulin, suggesting the inulin was significantly smaller than SAX in molecular size.

### Carbohydrate structure governed fermentation rate and metabolic outputs

We used inulin and SAX as substrates for *in vitro* sequential batch cultivation experiments, matched with monosaccharide controls mimicking glycosyl residue stoichiometry in the cognate polymer (fructose and SAX sugars, respectively) over 7 sequential passages (1/day). This sequential batch culture regime imposed very high dilution rates (10^-14^ dilution over 7 days), as in experimental evolution experiments conducted by Lenski and coworkers (18). Fermentation was vigorous for all samples in the first passage, as measured by both gas production and pH. Dilution initially reduced gas production (Figure 1A) in the second passage (day 2); monosaccharide control and inulin cultures stabilized in gas production resulted after day 3, whereas gas production from SAX remained low throughout the experiment. Terminal pH increased until day 3 for all conditions, after which it stabilized for SAX, SAX sugars, and inulin; terminal pH of fructose cultures continued to decrease until day 5 (Figure 1B).

**Figure 1.**
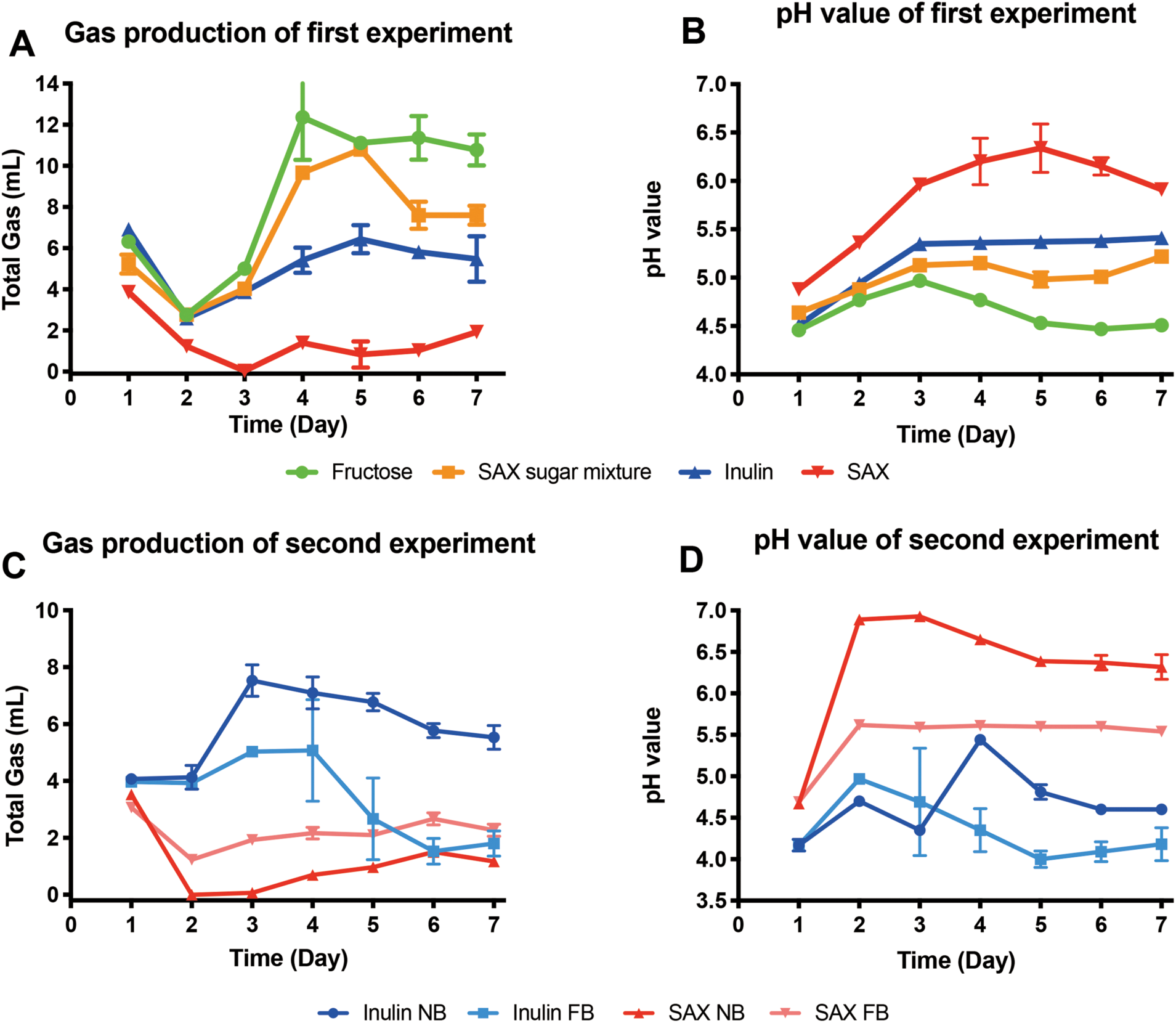
Total gas and acid production from sequential batch cultures of monomeric (fructose, SAX sugar control) and polymeric (inulin, SAX) carbon sources in unfortified media. Gas production as overpressure (A) and pH (B) were measured after each daily passage. Total gas (C) and acid production (D) from sequential batch cultures on polymeric (inulin, SAX) carbon sources in normal and fortified media were also recorded each day in the nutrient supplementation experiment. Error bars depict standard error of the mean.

The SCFAs acetate, propionate, and butyrate are regarded as the major terminal products of carbohydrate fermentation by human gut microbial communities. SCFA profiles produced from fructose, SAX sugars, inulin, and SAX are illustrated in Figure 5. Acetate production from all carbohydrates was largely steady or increased only slowly after day 2, except for SAX sugars. The highest acetate concentration from final cultures was observed for SAX sugars (21.1 mM), followed by SAX (15.0 mM), inulin (12.0 mM) and fructose (11.2 mM) (Figure 2). Only SAX elicited appreciable propionate and butyrate over sequential cultures in fortified buffered medium (FB), which contained amino acids and vitamins. Inulin has been reported to produce large amounts of SCFAs, especially butyrate (19), and did so in the first culture. However, over sequential passage inulin-fermenting consortia produced relatively lower total SCFAs (12.9 mM) and only trace amount of propionate (0.7 mM) and butyrate (0.2 mM). This result was consistent with relatively more of the carbon being released as CO_2_ and/or other, unmeasured organic acids rather than SCFAs (which is consistent with low terminal pH results from these cultures). Comparatively, fermentation of SAX produced much larger amounts of propionate (5.5 mM) and butyrate (2.2 mM) in FB at day 7; the normalized molar ratios of acetate, propionate, and butyrate in the final cultures were 66:24:10 (SAX), 98:1:1 (SAX sugars), 93:5:2 (inulin) and 94:5:1 (fructose) (Figure 2).

**Figure 2.**
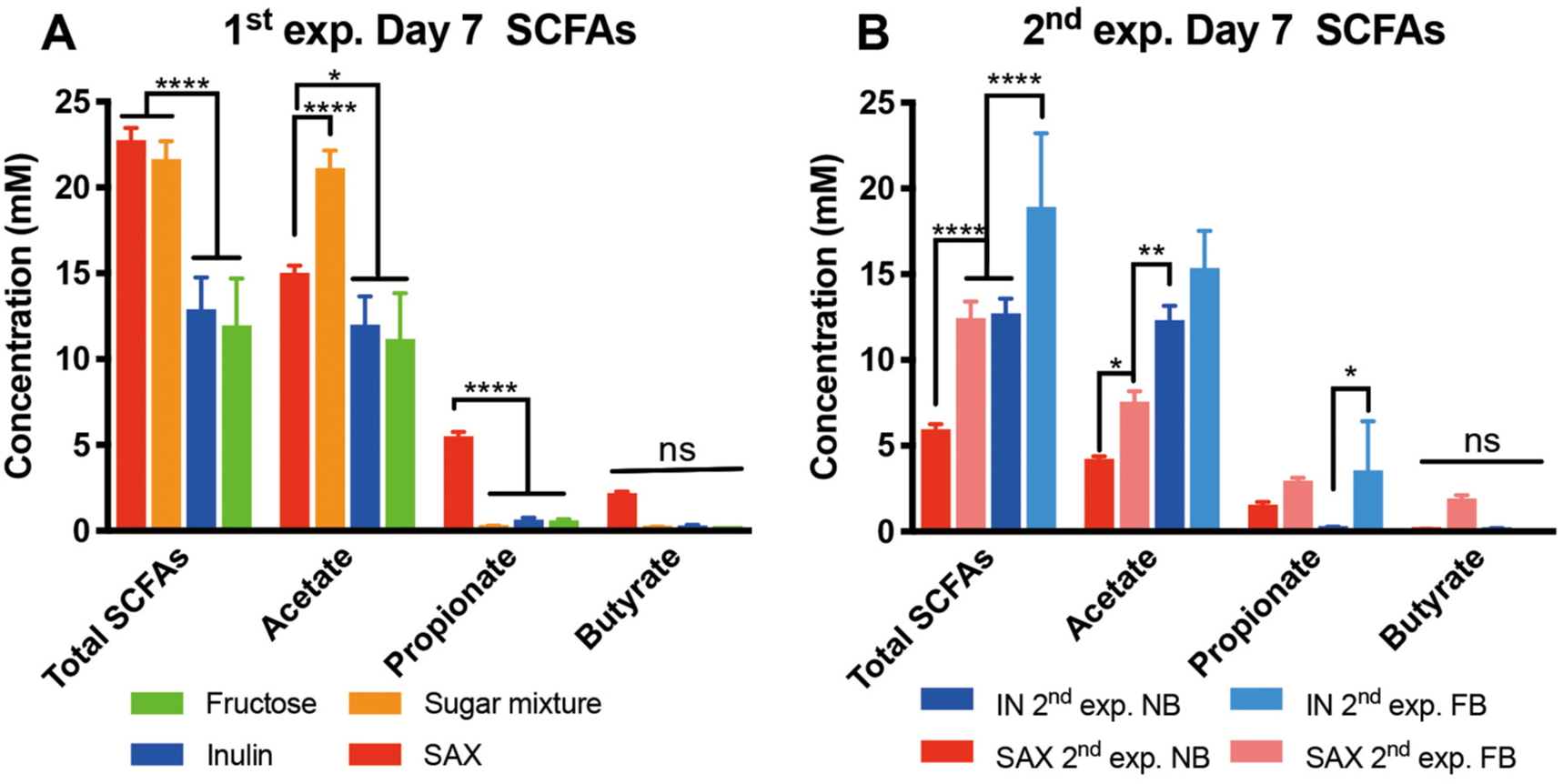
Comparison of total SCFA, acetate, propionate and butyrate production in day 7 cultures. (A) The SCFAs among SAX-, SAX sugar control-, inulin- and fructose-consuming cultures from the first experiment in NB. (B) SCFAs comparison plot of the second experiment in which SAX and inulin were fermented either in unfortified or fortified media. Error bars depict standard error of the mean. Statistically significant differences are calculated by Tukey’s multiple comparisons test with p < 0.05. Symbol style: nonsignificant (ns), 0.0332 (*), 0.0021 (**), 0.0002(***), <0.0001(****).

### Carbohydrate polymers maintained diverse and stable consortia

To determine whether carbohydrate structure influences the diversity of fermenting consortia, we sequenced the V4-V5 regions (20) of the 16S rRNA gene at each passage. Microbial relative abundances are plotted for one of the three replicate lineages for each condition and experiment in Figure 4. Across both experiments we describe here, microbial succession was very similar across the three lineages of each condition, suggesting carbohydrate structure-based selection exerts strongly deterministic forces on community assembly (Supplementary Figure 2). Interestingly, the first passage from fecal inocula were very similar in structure even across experiments (as seen by Bray-Curtis dissimilarity; Figure 3), with dominant populations of *Bacteroides* spp., diverse OTUs representing phylum *Firmicutes* and classified within families *Clostridiaceae, Lachnospiraceae, Enterococcaceae, Streptococcaceae,* and two OTUs within class *Negativicutes* (identified as *Phascolarctobacterium faecium* and *Veillonella* sp.; Figure 4A). In the second passage and thereafter, significant differences in succession emerged among conditions. In fermentation of inulin, fructose, and SAX sugar controls, cultures were strongly dominated by a bloom of OTU3 (*Escherichia* sp.), which was succeeded by OTU1 (*Klebsiella pneumoniae*) and remained dominant thereafter. Monosaccharide controls ended in near-monocultures of OTU1 (99.2% for fructose and 97.8% for SAX sugars). Inulin cultures also maintained small populations of *Clostridium ramosum* (OTU8, 5.7%) and *B. ovatus* (OTU2, 3.5%). Conversely, throughout the experiment SAX sustained a diverse community anchored by *Bacteroides ovatus* (OTU2, 63.7%), but with significant representation of OTUs classified within *Firmicutes* (especially *Eisenbergiella tayi* (OTU5, 13.6%) and *Hungatella effluvia* (OTU16, 3.5%)).

**Figure 3.**
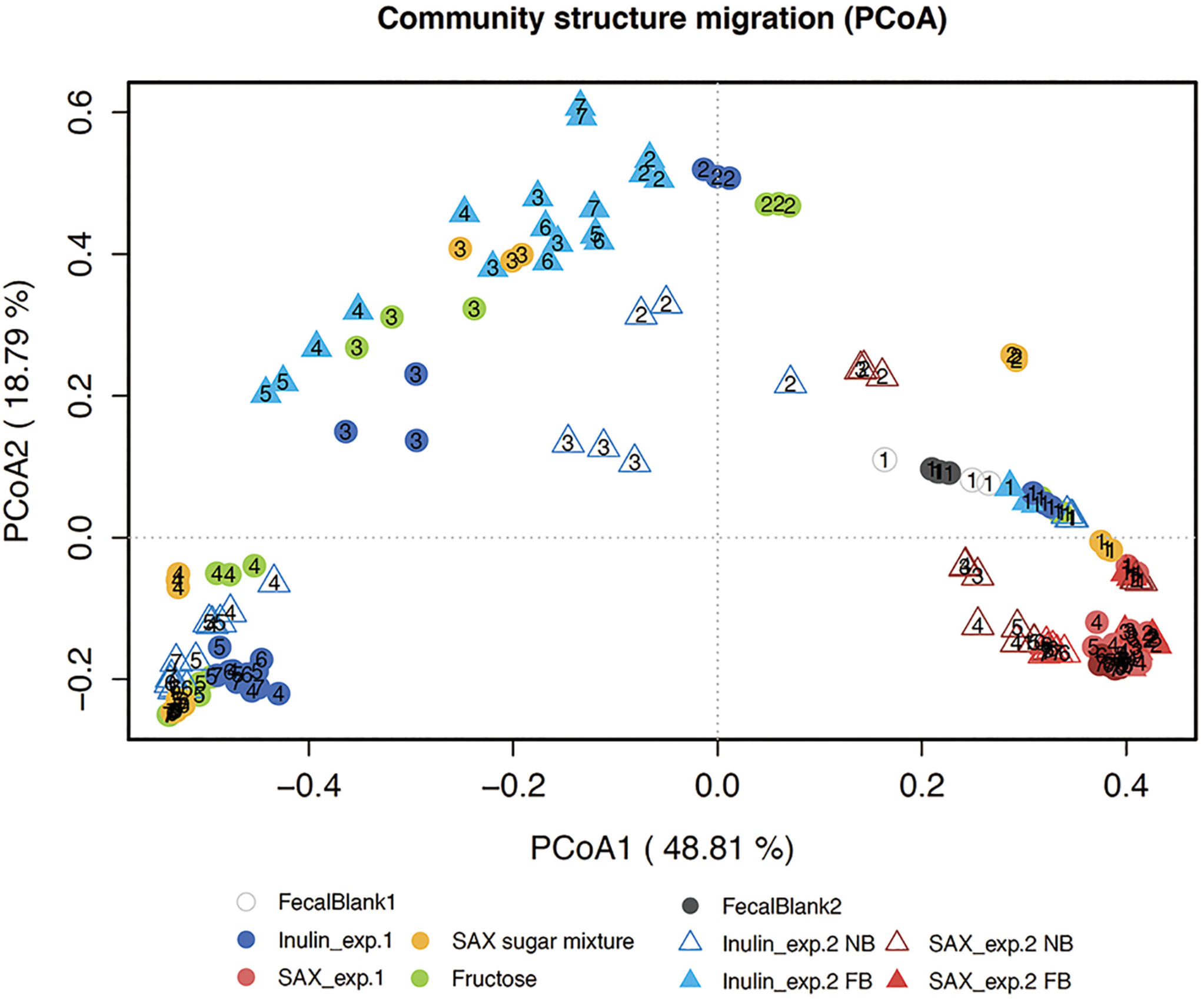
Principal coordinate analysis (PCoA) plot of Bray-Curtis dissimilaritiy across all lineages of both experiments. Numbers in the symbols indicate the passage day. SAX-consuming consortia clustered in the lower right corner across both experiments, indicating similar community structure regardless of medium condition. Inulin-consuming consortia in normal media formed a distinct cluster, which was close to the fructose and SAX sugars control samples. In contrast, communities consuming inulin in fortified media displayed an alternate succession trajectory and resulted in distinct community structure.

**Figure 4.**
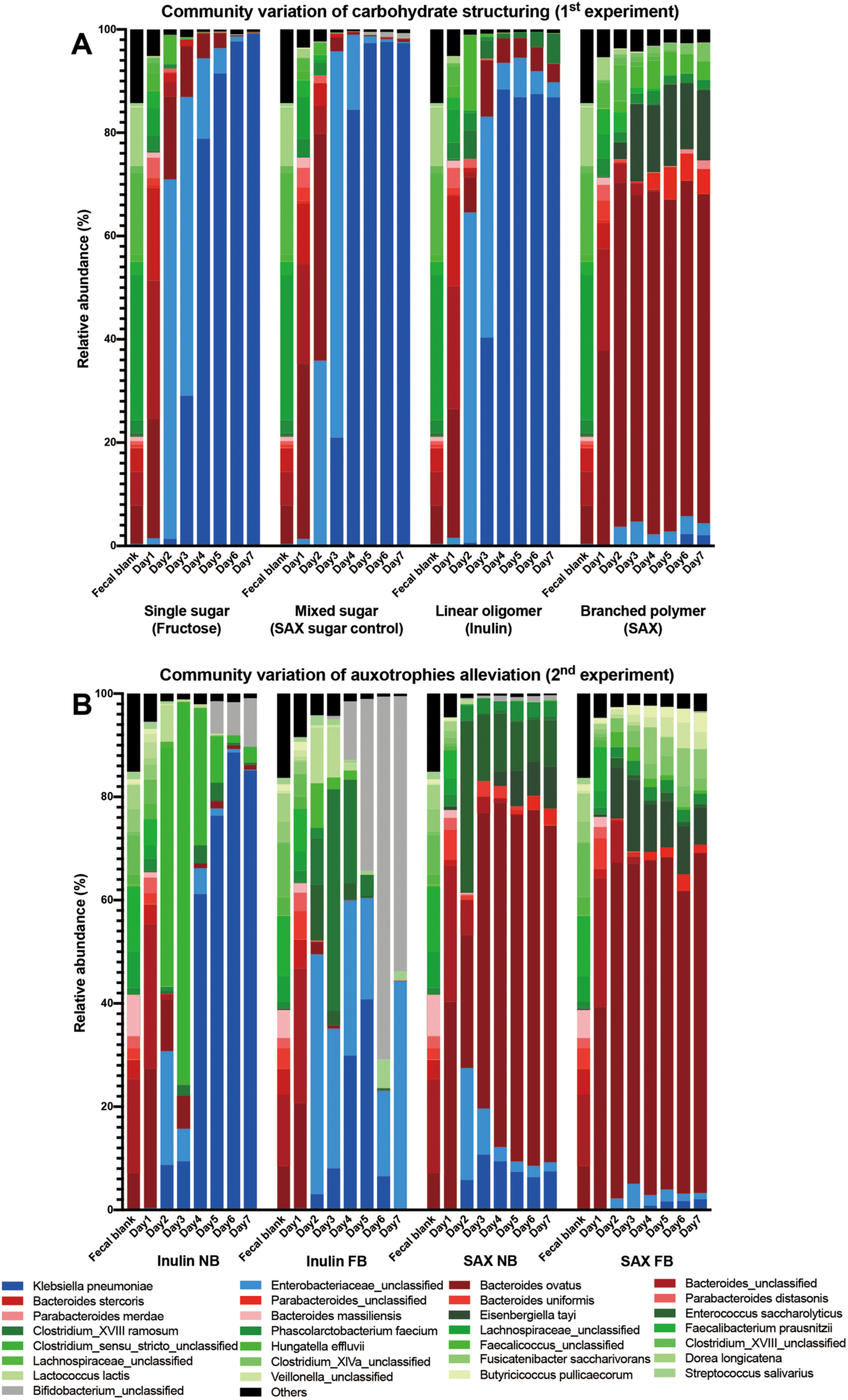
Microbial community structure across sequential passages as revealed by 16S rRNA gene amplicon sequencing. Daily changes of community structures from one donor’s lineage over 7 days are shown in (A) sequential cultures on inulin, SAX, and their respective monosaccharide controls (first experiment) and (B) the effect of fortification on SAX- and inulin-fermenting consortia (second experiment). Each shade represents a distinct OTU, which are colored by phylum: *Proteobacteria* (blue), *Bacteroidetes* (red), *Firmicutes* (green) and *Actinobacteria* (gray). Rare OTUs (those below 0.2% relative abundance) are grouped in Others (black).

**Figure 5.**
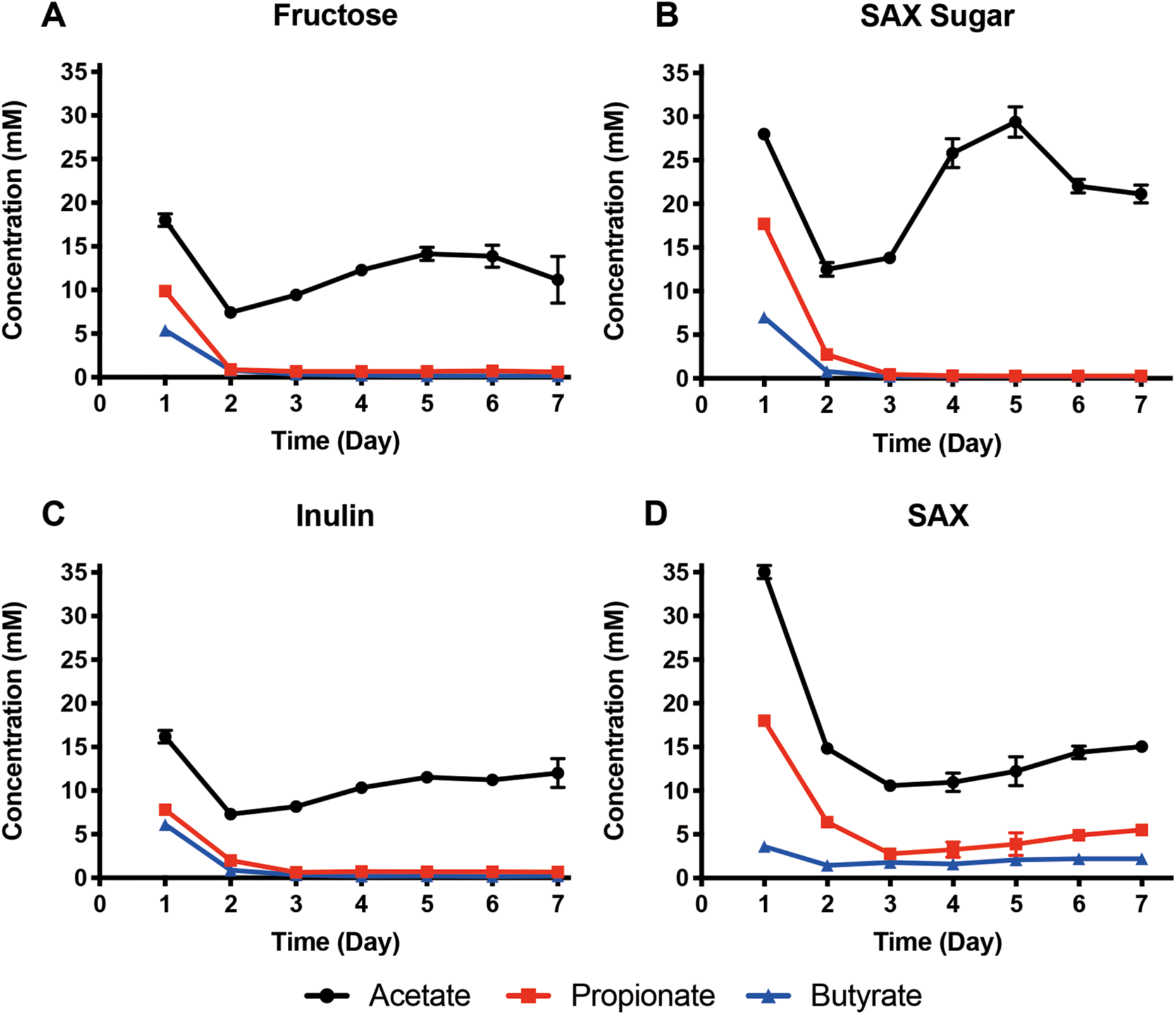
Short-chain fatty acid (SCFA) production as a function of carbohydrate complexity, including polymers (inulin, SAX) and their corresponding monosaccharide controls (fructose (A), SAX sugar control (B), inulin (C) and SAX (D)). SCFAs were measured at each daily passage by gas chromatography. Error bars depict standard error of the mean.

### Alleviation of obligate interactions imposed by auxotrophy drastically altered succession of inulin- but not SAX-fermenting lineages

To test the hypothesis that carbohydrate polymer structure, rather than other metabolic interactions among members (imposed by their inability to biosynthesize required components) governed composition, we performed a similar sequential batch fermentation experiment using inulin and SAX as sole carbon sources but with and without supplementation of amino acids and vitamins. Fecal inocula were derived from the same donor as the first experiment, collected fresh one month later. In contrast to the initial experiment, gas production from inulin did not decrease at passage 2 and 3, however SAX fermentations displayed similar dynamics across experiments (Figure 1C). Gas production from inulin was inverse to acid production, decreasing with concurrent decreases in pH slightly for inulin-fermenting consortia in normal buffered medium (NB) and much more strongly in FB (Figure 1D). In both cases, this shift was accompanied by transitions from day 3 and 4 cultures dominated by clostridia and enterobacteria to increasing bifidobacteria (which ferment largely to acetate and lactate) (21), though the magnitude was much larger with fortification (see below). Fortification significantly increased both gas and acid production from SAX.

SCFA production data was consistent within inulin samples, which was acetogenic with and without fortification; propionate and butyrate concentrations were low or undetectable after a peak of butyrogenesis in NB on day 3 (Figure 6). Production of SCFAs from SAX in NB was substantially smaller than in the first experiment; small amounts of acetate (∼5 mM) and propionate (∼1.5 mM) were detected but butyrate was vanishingly small (∼0.1 mM). With fortification, relatively stable production of acetate (∼10 mM), propionate (∼3 mM), and butyrate (∼2 mM) were observed after passage 2 (Figure 6). These data, coupled with the gas production data, suggested that the utilization of SAX was rate-limited in normal medium and alleviated with fortification. Though SAX is regarded as a highly propiogenic fiber in fecal fermentation (22), our data also suggested sustained butyrate production with fortification. We detected trace amounts of other fermentation products including lactate and ethanol (Supplementary Table 1) in day 7 SAX-fermenting culture supernatants; they did not approach SCFA concentrations and did not vary with medium fortification.

**Figure 6.**
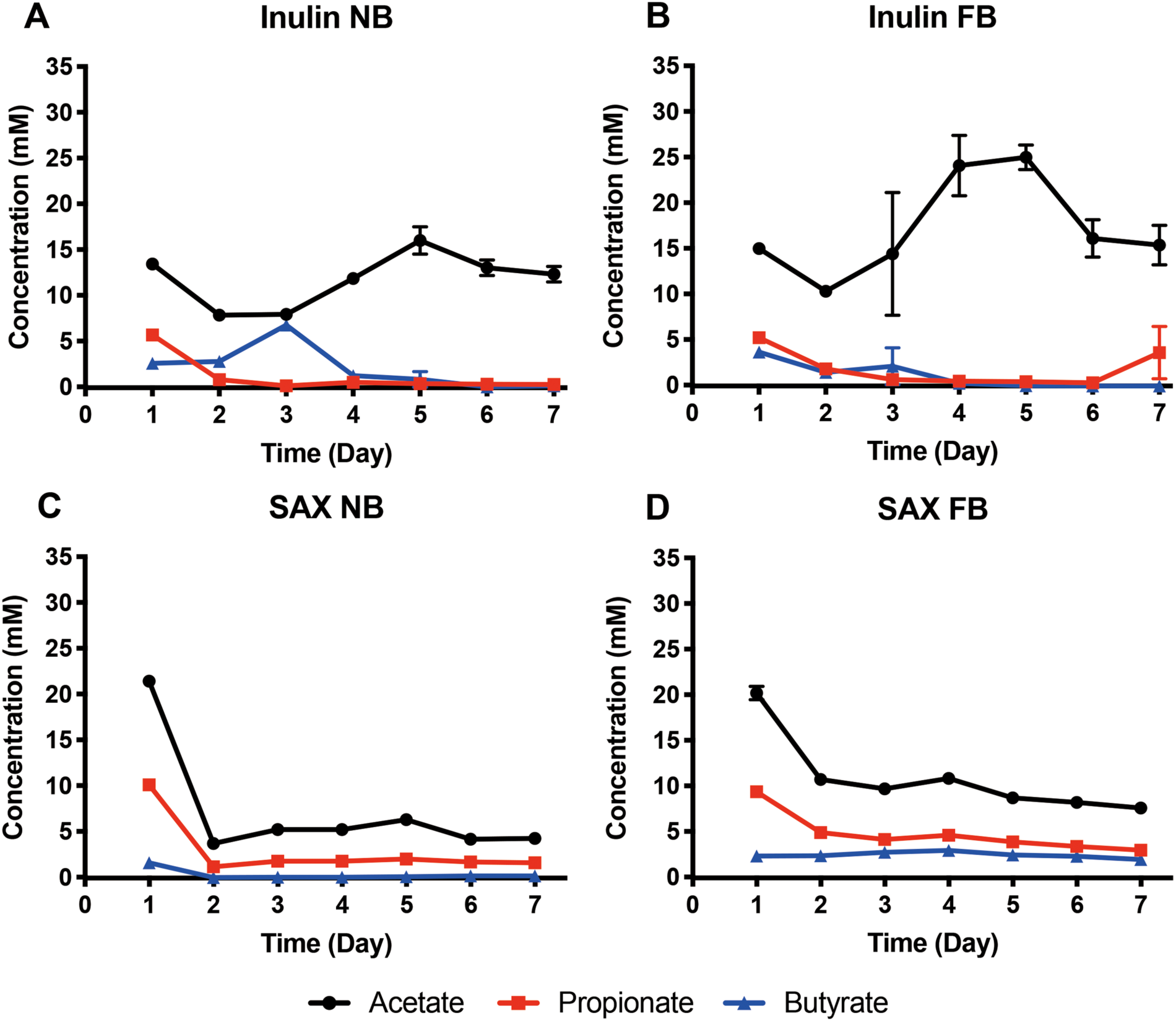
Short-chain fatty acid (SCFA) production of inulin- and SAX-consuming cultures with and without medium fortification with amino acids and vitamins. SCFAs were measured at each daily passage by gas chromatography. Error bars depict standard error of the mean.

Alleviation of metabolic interdependencies imposed by auxotrophies exerted a much stronger impact upon inulin-fermenting than SAX-fermenting community structure. β-diversity analyses revealed clustering of SAX-consuming consortia across experiments and media conditions (Figure 3), driven most strongly by similarly dominant OTU2 *B. ovatus* populations (Figure 4A,B). However, inulin-fermenting consortia in NB were dominated by OTU1, with smaller populations of clostridia (OTU8, *C. ramosum* in the first and OTU14 *Clostridium* sp. in the second experiment). Both the first and second experiment in NB-inulin ended in cultures dominated by OTU1. However, fortification resulted in very different succession trajectories; communities initially dominated by *Bacteroides* spp. (OTU2 and OTU4) then yielded to *Clostridium* sp. (OTU14 in NB and OTU8 with fortification). Thereafter, OTU1 became dominant in NB, with minor populations of OTU2 and OTU8. With fortification, *Bifidobacterium* sp. OTU6 became dominant with a sizeable population of OTU3. Thus, fortification governed both bifidobacterial abundances at the conclusion of the experiment and the identity of the dominant enterobacterial species, resulting in very different final communities.

In contrast, SAX-fermenting consortia displayed remarkably similar successional trajectories across experiments and media conditions. Besides the aforementioned dominance of OTU2 across all SAX cultures, many other taxa were shared (e.g. OTU5 *Eisenbergiella tayi*, OTU9 *Phascolarctobacterium faecium*), though at different relative abundances. Interestingly, final consortia from the first experiment’s NB were more similar in structure to FB in the second, though only modestly so. One example is in the proteobacterial component; in the first experiment and in fortified cultures, these populations were uniformly small throughout succession and ended with approximately equal fractions of OTU1 and OTU3. However, in the second experiment’s NB condition, a proteobacterial bloom (especially OTU3) occurred on passage 2, sharply declining thereafter and stabilizing with larger OTU1 populations. Further, *Clostridium* sp. OTU14 displayed a concurrent bloom only in NB. Similarity between the first experiment’s SAX NB consortia and second’s FB consortia suggests that similar taxa were present in the inoculum at low abundances, being retained over a one-month time interval in the donor’s microbiome.

### Carbohydrate structural complexity governed consortial diversity

Carbohydrate structural complexity most strongly governed maintained diversity across sequential passages, but diversity was also influenced by medium fortification. As expected, carbohydrate polymer fermentation sustained consortia of significantly higher α-diversity than monosaccharide controls (Figure 7, panels A, C, E, G) over sequential passages. SAX-consuming consortia stabilized in the first experiment at ∼19 OTUs of appreciable abundance (> 0.1%; Fig. 8A), which was far higher than other lineages (including inulin) and was reflected in Shannon and inverse Simpson metrics (Fig. 8C, G). These data suggested that structured carbohydrates retained greater microbial diversity over sequential passages, and the magnitude of the sustained diversity was related to the polymer’s complexity.

**Figure 7.**
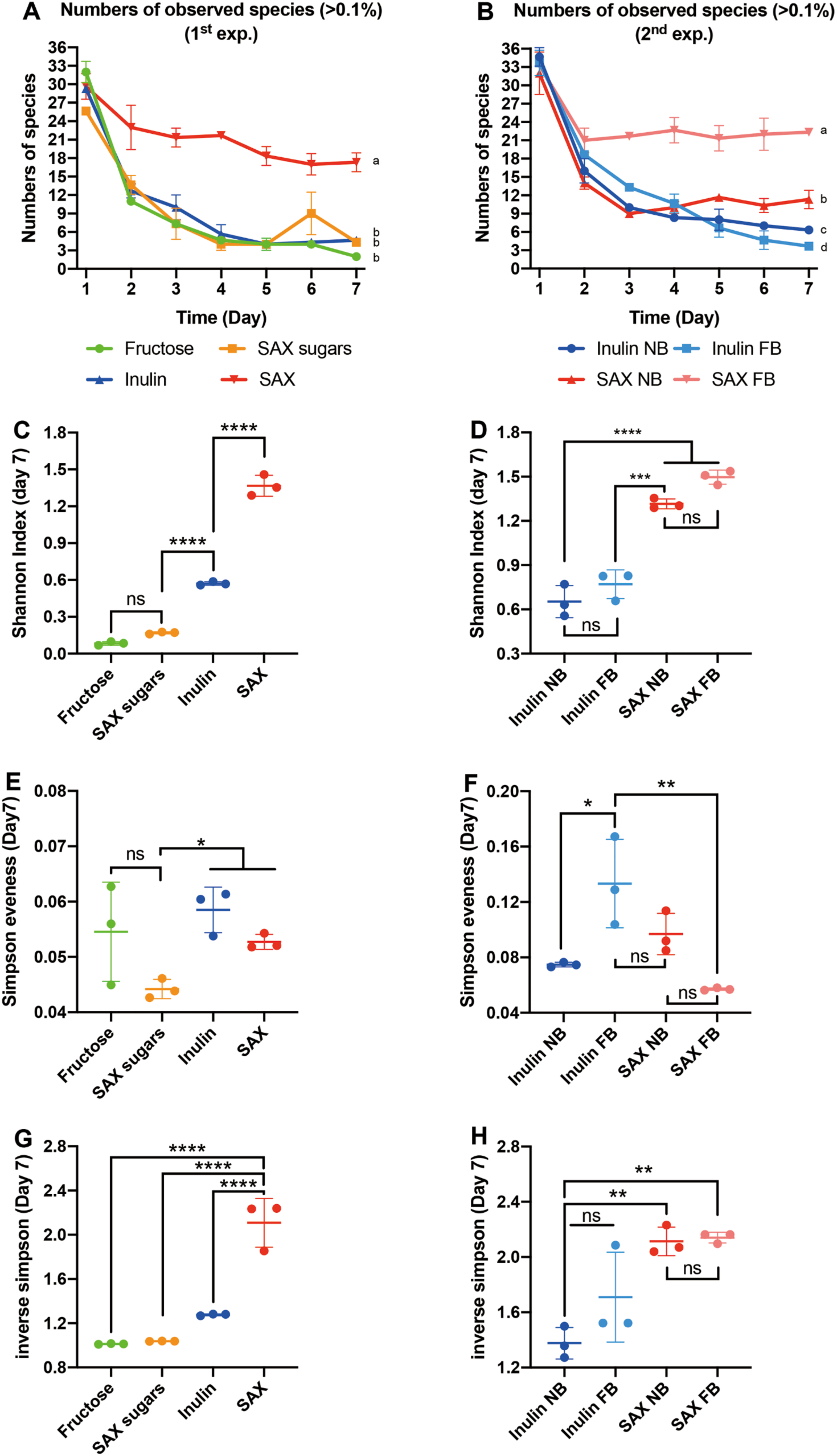
Numbers of observed OTUs (> 0.1% relative abundance across each sample) at each passage in the first experiment (A) and second experiment (B). Alpha diversity metrics (Shannon index (C,D), Simpson evenness (E,F) and inverse Simpson index (G,H) of day 7 communities are plotted for each experiment (first experiment: A, C, E, G, second experiment: B, D, F, H). Identical letters in panels A and B indicate nonsignificantly different values. Statistically significant differences are calculated by Tukey’s multiple comparisons test with p < 0.05. Symbol style: nonsignificant (ns), 0.0332 (*), 0.0021 (**), 0.0002(***), <0.0001(****).

Fortification increased the number of apparent species (OTUs greater than 0.1% in abundance, which corresponded to an average of ∼2.6 reads) in SAX-fermenting consortia but did not significantly influence Shannon, inverse Simpson, and Simpson evenness (Figure 7 D, F & H). However, the opposite result was observed for fortification of inulin-fermenting cultures; fortified inulin consortia displayed fewer species observed but significantly increased evenness, driven mostly by relatively even abundances of OTU3 and OTU6 in fortified cultures. Taken together, these data suggest that complex polysaccharides sustain diverse fermenting consortia over sequential dilutions in a manner related to the structural complexity of the polysaccharide, and that the extent to which carbohydrate structure drives final community composition, as opposed to other metabolic interdependencies, differs with polymer structural complexity.

## DISCUSSION

The sequential batch fermentation approach, also called consecutive batch culture, has been used for decades to understand the associations of microbiota with the polysaccharides they hydrolyze, but to the best of our knowledge this approach has not been widely employed to identify linkages between human gut microbiota and the dietary carbohydrates they consume. For example, Gascoyne and Theodorou passaged rumen microbiota ten times through a medium containing soluble carbohydrates and rye grass hay, finding that the addition of monosaccharides into hay fermentations changed the molar ratio of SCFAs and the fraction of biomass digested (23). Cheng *et al.* passaged rumen digesta nine times through a media containing 1% barley straw and found that the fiber-degrading consortia were robust and permitted transfer for up to 63 days (24). However, previous studies before the advent of culture-independent assays were hampered in their ability to measure the effect of carbohydrate polymer structure on maintenance of diversity. We employed a similar approach with amplicon sequencing to identify the diversity of human gut microorganisms selected by model carbohydrate structures under *in vitro* conditions.

Our results support the hypothesis that linkage diversity allows microbial division of labor in consumption of structured carbohydrates, which in turn permits cooperative consumption of these carbohydrates and maintains microbial diversity even under high dilution pressure. Further, they suggest that microbial succession on carbohydrates is influenced by the availability of other nutrients, which disrupts obligate metabolic interactions imposed, presumably, by auxotrophy. It should be noted, however, that trace nutrients may subtly alter fitness landscapes in other ways beyond binary growth/no growth phenotypes imposed by auxotrophy; nutrient availability may, for example, also exert impacts through alterations in gene regulation, enzyme activity, or metabolic fluxes.

Recently, Chung et al. employed continuous flow fermenters (bioreactors) to simulate the human colonic environment for 20 days to investigate the relationship between microbial diversity and structural complexity of dietary fibers (25). Inulin and arabinoxylan-oligosaccharides (AXOS) were used as substrates in their study, although the continuous cultivation method employed imposed much less dilution pressure (one medium exchange per day) than our experiment. Their results showed dominance by members of family *Bacteroidaceae* across all carbon sources, suggesting strong priority effects strongly influence community structure at low dilution (similarly to the first passages in our experiment). However, though they did not perform detailed carbohydrate structure analyses, they also observed that dietary fibers with likely-greater structural complexity sustained greater microbial diversity in the reactors, which concurs with our findings. Together, their data and ours suggest that relationships between carbohydrate structural complexity and sustained diversity may be a fundamental property of carbohydrate-microbiome interactions.

In our study, the fructose control was the simplest carbon source that can be widely transported and utilized by most anaerobic bacteria. With the mixture of two pentoses and three hexoses, the SAX-simulating sugar mixture retained slightly higher diversity compared with the fructose culture at day 3 (Figure 4), but finally stabilized with similar consortia after day 4, indicating that monosaccharide mixes have only a weak ability to maintain diversity. Inulins are fructan-type storage carbohydrates from plants such as chicory, onions, and asparagus (26), in which one terminal glucosyl unit is linked with an α-1,2 bond to fructose (as in sucrose) and is extended into chains of β-2,1-linked fructose residues (DP = 2–60). In this study, inulin was used as an intermediate-complexity carbon source since it contains only two types of glycosidic linkages in its linear chain structure. Inulin day 7 consortia showed a significantly higher Shannon index than those fermenting monosaccharides, indicating that even simple carbohydrate structuring supports maintenance of microbial diversity. Arabinoxylans are hemicelluloses typically composed of a β-1,4-linked xylose backbone, substituted at the 2- and 3-carbon positions with arabinose branches. The branches can be disaccharides or even trisaccharides, and was employed as a model complex carbon source structure in this study. The molecular size of AXs can range from 300 kDa to 2 MDa (27). Compared with inulin, AXs have more than 10 different linkage types that require multiple CAZymes to fully decompose. Therefore, day 7 SAX-consuming consortia exhibited very different community structures (*Bacteroides* spp.-dominant) with the highest diversity among four carbon sources.

It is notable that congruence was observed across two experiments from the same donor’s microbiota separated by a month and in by very high similarities among three independent replicate lineages for each carbohydrate, which stabilized with microbial diversities that related to structural complexity (Figure 4, Supplementary Figure 2). As we observed the same OTUs across both experiments, our data suggest that the same V4-5 ribotypes responsive to SAX and inulin were present in this individual’s microbiome across this time frame, despite an uncontrolled diet. However, some of the taxa observed in the first (unfortified) SAX-consuming consortia required alleviation of auxotrophy in the second to become dominant. These data suggest the possibility that the initial availability of “public good” nutrients (28) like amino acids and vitamins (or natural networks of producers of these nutrients that influence exchange) may have varied among experiments, driving differences in community succession. Further, it suggests the possibility that variations in the initial concentrations of gut micronutrients or natural assemblages of microbiota that impact community nutrient exchange may impact the activity and abundances of microbiota in *in vitro* fermentations.

If initial nutrient conditions substantially vary in *in vitro* fecal fermentations, even from the same donor, these differences may influence the outcomes of batch fermentations attempting to link carbohydrates with fermenting microbiota. This suggests tighter control of initial nutrient conditions in fecal fermentations is likely warranted to identify relationships between carbohydrate structure and fermenting microbiota. Increases in microbial abundances may arise due to extensive metabolic interactions in diverse communities that are second- or third-order to consumption of the carbohydrate substrate. If true, this suggests that single batch cultures from fecal inocula may mask the organisms able to grow most rapidly on a certain carbohydrate. Together, our data suggest substantial context-dependence in carbohydrate utilization by gut communities, which declines with increasing structure.

This context-dependence may help explain conflicting results in *in vitro* and *in vivo* experiments linking resistant carbohydrates to the microbiota they selectively favor (central to the definition of a prebiotic (29). Inulin and fructooligosaccharides (FOS) have been extensively studied for their bifidogenic effect. In human trials, increases in bifidobacterial populations after consumption of inulin or FOS have been regularly observed (30, 31). Nevertheless, contradictory results of the inulin bifidogenic effect have also been reported. Bettler and Euler supplemented infant food with FOS for 212 infant participants and found no significant change of bifidobacterial count over 12 weeks of study (32). Calame *et al.* provided 54 volunteers with inulin for 4 weeks but observed no bifidogenic effect (33).

Investigations have demonstrated that inulin interactions with gut microbiota may not be highly specific to promoting growth of *Bifidobacterium* spp. in mixed communities. Scott *et al.* cultured both short-chain and long-chain inulin samples with dominant human colonic butyrate producers and *Bifidobacteria*, finding that all 15 tested strains including *Faecalibacterium*, *Roseburia*, *Eubacterium*, *Anaerostipes*, *Bifidobacterium* and *Bacteroides spp.* were able to consume short-chain inulin (DP 2-8), while only *Roseburia inulinivorans* was able to propagate on long-chain inulin (DP 25) (34). Sheridan *et al.* analyzed genomes of *Rosburia* spp. and *Eubacterium* spp. and found *Roseburia inulinivorans* and *Agathobacter rectalis* (previously *Eubacterium rectale*) strains also exhibited genomic evidence of inulin utilization genes and, correspondingly, strong growth on inulin (35). These data suggest that small alterations in carbohydrate structure may alter which organisms are able consume a carbohydrate, and, specifically, that the organisms best able to grow on inulins may depend upon community context (i.e., population sizes of competing organisms). Our data further suggests that concentrations of other nutrients may also shape community responses to inulins.

With respect to members of family *Enterobacteriaceae*, *Klebsiella pneumoniae* has been previously observed to grow on fructooligosaccharides, which differ from inulins only in the terminal glucose residue (36). Reports are few on the utilization of inulin and FOS by *Escherichia* spp. However, Hidaka *et al.* reported that *K. pneumoniae* was able to consume FOS, but *E. coli* did not (37). Hartemink *et al.* observed the growth of *E. coli* and *K. pneumoniae* in media containing FOS (38). Schouler *et al.* identified a pathogenic *E. coli* strain BEN2908 containing a *fos* locus that utilized kestose and nystose (inulin of DP = 4) (39). Although most *in vivo* studies suggest that inulin and FOS feeding are associated with inhibition of pathogenic bacteria in the colon, our data and these observations suggest the inhibition phenomenon may actually originate from the success of competing microbial consortia rather than the selectivity of the substrate for bifidobacteria and lactobacilli.

Members of *Bacteroidetes* are known to possess the ability to utilize complex dietary carbohydrates and also encode CAZymes for cleavage and the intake of oligosaccharide substrates (40). Human gut bacteroides, including the strains of *B. eggerthii*, *B. cellulosilyticus*, *B. intestinalis*, *B. ovatus* and *B. xylanisolvens,* are generally able to utilize xylan (41). *B. ovatus*, as the dominant OTU in the final SAX consortia, have surface endoxylanases that are able to break down the xylan backbone extracellularly (42). Subsequently, the released xylan fragments (sometimes with arabinose substituents) are imported through the outer membrane via surface carbohydrate-binding proteins and are debranched in the *B. ovatus* periplasmic region. However, extracellular hydrolysis and import is not perfectly efficient for *B. ovatus*; the released oligosaccharides can support the growth of species that do not have xylan-consuming ability (43). Little information exists on microbial populations that are associated with *B. ovatus* xylan consumption. Our sequential passage approach suggested possible interactions among *B. ovatus*, *Eisenbergiella tayi* and a *Clostridium* XIVa species (Figure 4B). *Eisenbergiella tayi* produces acetate, butyrate and lactate as its major metabolic end products (44). Members of *Clostridium* XIVa are known as major butyrate producers in the colon, which may have a beneficial effect on gastrointestinal health (45). Increased abundances in both of these taxa in fortified versus unfortified media may explain the differential butyrate concentrations among SAX-consuming consortia.

Modulating human gut microbiota using dietary carbohydrates has been proposed as a possible approach to improve human health. The composition of gut microbiota is affected by the carbohydrate degradation process and the interactions of key carbohydrate-degrading microorganisms. Although there are variations in gut microbiota structures among individuals, the loss of community diversity has been associated with multiple health disorders such as obesity and Crohn’s disease (6). Our *in vitro* fermentation data suggest the hypothesis that complex carbohydrate structure may be a potential mechanism to maintain diverse human gut microbiota *in vivo*.

## MATERIALS and METHODS

### Carbohydrate substrates

Inulin was purchased from a commercial chemical supplier (Alfa Aesar, Haverhill, MA); the inulin sample contains fructan polymers up to degree of polymerization (DP) of 25 according to the manufacturer’s specification sheet. SAX was isolated in the lab from sorghum bran using alkali extraction followed by an ethanol precipitation method as described previously (46). Sugar controls including fructose, xylose, arabinose, mannose, galactose and glucose were purchased from Sigma-Aldrich Inc (St. Louis, MO).

### Polysaccharide composition and structure

The neutral monosaccharide compositions of inulin and SAX were determined using the acid hydrolysis and volatile alditol acetate derivatization methods as previously described (47, 48). Briefly, inulin polymers were hydrolyzed in 2M trifluoracetic acid (TFA) at 50°C for 30 min; sugar concentrations in hydrolyzate were quantified by HPLC (49). SAX was hydrolyzed in 2M TFA at 121°C for 90 min; the released sugars were reduced by NaBD_4_ and were converted into volatile alditol acetates by reacting with acetic anhydride at 100°C for 2.5 h. The alditol acetate derivatives were quantified via gas-chromatography-mass spectrometry (12).

Molecular size distribution of inulin and SAX was analyzed by high performance size-exclusion chromatography with refractive index detectors (Wyatt Technology Corporation, Santa Barbara, CA) as previously described (46). Briefly, sample solutions were prepared in DI water (0.1% w/v), prefiltered through a syringe membrane (1.5 μm) and injected (100 μL) into S500HR column (Amersham Biosciences, Piscataway, NJ). Data were collected after 130 minutes elution and RI signals were normalized by Origin Pro 9.1 (OriginLab Corporation, Northampton, MA).

### *In vitro* sequential fermentation

Fermentations were performed in an anaerobic chamber (Coy Laboratory Products Inc., Great Lake, MI) supplied with an 90% N_2_, 5% CO_2_, and 5% H_2_ atmosphere. Sodium phosphate buffer (10 mM) was prepared and autoclaved at 121°C for 20 min. The base media (hereafter referred to as normal buffered (NB) medium) contained (per liter) 0.47 g NaCl, 0.45 g KCl, 0.40 g urea, 0.10 g Na_2_SO_4_, 0.001 g resazurin, 0.865 g Na_2_HPO_4_ and 0.468 g NaH_2_PO_4_ and was autoclaved (121°C for 20 min). Heat sensitive compounds, including 0.0728 g CaCl_2_, 0.1 g MgCl_2_, 1 mL of 1000X P1 metal solution, and cysteine hydrochloride (0.25 g/L) were sterilized by 0.22 μm filtration and added; the media was then reduced in anaerobic chamber overnight. To alleviate auxotrophies via exogenous nutrients, media were also fortified with both 200 µM amino acids (10 μM of each) and 1% (v/v) ATCC vitamin supplement (Hampton, NH). This medium is hereafter referred to as fortified buffered (FB) medium.

The protocols involved for handling human fecal samples were reviewed and approved by Purdue University’s Institutional Review Board (IRB protocol #1701018645). Human fecal samples were acquired from a single healthy donor who had not received antibiotic treatment in the prior three months. Samples were sealed in 50 mL Falcon tubes, stored on ice and followed by a rapid transfer into an anaerobic chamber. To prepare the inoculum, fecal samples were anoxically homogenized in sodium phosphate buffer at a ratio of 1:20 (w/w). The fecal slurry was filtered through four layers of sterile cheesecloth and used as inocula within 2 hrs. Fresh fecal inocula used in the two described experiments were collected from the same donor at a one-month interval. The overall research scheme is illustrated in Supplementary Figure 3.

Sequential batch cultivations of inulin and SAX-consuming consortia were conducted in 25 mL autoclaved Balch-type culture tubes (Chemglass Life Sciences, Vineland, NJ) in triplicate lineages (5 mL each, with total carbohydrate of 1% w/v). The tubes were sealed with rubber stoppers (Chemglass Life Sciences, Vineland, NJ) and aluminum seals (Chemglass Life Sciences, Vineland, NJ), and incubated in an Innova 42 shaking incubator (New Brunswick Scientific, Edison, NJ) at 37°C and 150 rpm for 24 hrs. For sequential passages, tubes were opened inside the anaerobic chamber to maintain low oxygen condition in each passage, diluted 1:100 into fresh media and fermented for another 24 h. The sequential cultivation experiment was continued for 7 days, and each substrate/culture condition was cultured in triplicate lineages that were not intermixed.

The gas production (as overpressure volume) of each culture tube was recorded for each passage by piercing the stopper with a needle and glass syringe. Two aliquots were collected from cultures at each passage for DNA extraction (1 mL) and short-chain fatty acid measurements (1 mL). These samples were stored in sterile Eppendorf tubes and immediately frozen at −80°C. The pH value was recorded from the remainder using a pH meter (Mettler Toledo, Columbus, OH).

### Short chain fatty acid (SCFA) and other metabolites analysis

SCFAs were quantitated as described previously (50). Briefly, cultures were centrifuged (13,000 rpm) for 10 minutes to remove cell debris, mixed 10:1 with an internal standard mixture containing 4-methylvaleric acid, phosphoric acid, and copper sulfate pentahydrate, and supernatants were transferred to 2 mL screw-thread autosampler vials. SCFA (acetate, propionate and butyrate) concentrations were quantified using an Agilent 7890A gas chromatograph (GC-FID 7890A, Santa Clara, CA) with a fused silica capillary column (Nukon SUPELCO No:40369-03A, Bellefonte, PA).

SAX consortia culture at day 7 were analyzed for formate, lactate, succinate and ethanol concentrations using a HPLC system (Waters Corporation, Milford, MA) that was equipped with an organic acid column (BioRad Aminex HPX-87) and a refractive detector (Model 2414, Waters Corporation, Milford, MA). The SAX culture (1 mL) was centrifuged at 13,000 rpm for 10 min and then the supernatant was filtered through a 0.2 µm membrane to acquire a cell-free solution. The sample injection volume was set at 30 µL, the column was operated at 50°C and was eluted using 0.005M H_2_SO_4_ at 0.6 mL/min, the run time was set at 25 minutes.

### DNA extraction

Microbial DNA was extracted using a modified phenol-chloroform method (51, 52). Briefly, 1 mL of fermentation sample stored for DNA extraction was incubated at 85°C for 5 min to inactivate native DNAses, cooled, and centrifuged at 13,000 rpm for 10 min. The resulting cell pellets were incubated with 500 µL lysozyme (Fisher Bioreagents, Pittsburgh, PA) solution (1 mg/mL) at 37°C for 45 min. 100 µL proteinase K (Fisher Bioreagents, Pittsburgh, PA) solution (0.2 mg/mL) was then added and further incubated at 56°C for 1 h. Lyasates were then extracted with 500 µL phenol/chloroform/isoamyl alcohol (Acros Organics, Morris Plains, NJ) solution (25:24:1, PCI) in a reinforced sterile microvial (330 TX) preloaded with 0.3 g of 0.1 mm zirconia/silica beads (1107910z, both BioSpec Products, Inc., Bartlesville, OK). The lysis tubes were subjected to bead beating in a FastPrep-24 homogenizer (MP Biomedicals, Santa Ana, CA) for 10 sec (6 m/s), which was followed by cooling on ice for 5 min. An additional 500 µL of PCI solution were added and vortexed for 15 sec. Samples were centrifuged (13,000 rpm, 10 min, 4°C), and the upper aqueous phase was transferred to new tubes and mixed with 1 mL of a chloroform/isoamyl alcohol solution (24:1) by vortex for 15 sec. Samples were centrifuged again (13,000 rpm, 10 min) and the upper aqueous phase was transferred to new tubes containing 100 µL of 3 M sodium acetate and precipitated with 1 mL of isopropanol. The solution was shaken gently, incubated at room temperature for 10 min, and centrifuged at 13,000 rpm for 30 min. Supernatants were decanted and the DNA pellets were washed with 500 µL 70% ethanol twice. The microbial DNA was then air-dried, resuspended in Tris-EDTA buffer (pH 8.0), and stored at −20°C.

### 16S rRNA amplicon sequencing, sequence processing, and community analysis

Regions of the 16S rRNA gene were amplified from microbial DNA and sequenced to quantify community composition as described previously (12). Briefly, primers 515-FB (GTGYCAGCMGCCGCGGTAA) and 926-R (CCGYCAATTYMTTTRAGTTT) (20) were used over 20 amplification cycles. The resulting amplicons were purified using an AxyPrep PCR Clean-up Kit (Axygen Inc., Tewksbury, MA) following the manufacturer’s instructions. Amplicons were then barcoded using the TruSeq dual-index approach for five cycles. The barcoded amplicons were purified as above, quantified using a Qubit dsDNA HS Assay Kit (Invitrogen, Carlsbad, CA) and pooled. Quality control for each pool was performed on an Agilent Bioanalyzer (Agilent, Santa Clara, CA) and sequenced on an Illumina MiSeq run with 2 x 250 cycles at the Purdue Genomics Core Facility. Sequences are associated with BioProject PRJNA432190 and publicly available as BioSamples SAMN08438641-SAMN08438684 of the National Center for Biotechnology Information’s Sequence Read Archive. Sequences were processed using mothur v.1.39.3 according to the MiSeq SOP (https://www.mothur.org/wiki/MiSeq_SOP) with previously described modifications (50). Groups were subsampled to 2,594 reads to normalize sampling effort.

## ACKNOWLEDGEMENTS

This work was supported, in part, through Hatch project IND011670 to S.R.L. and through institutional funds provided by the Purdue University Departments of Food Science and Nutrition Science. The authors thank Dr. Anton Terekhov for his assistance with carbohydrate analyses.

---

Supplementary Figure 1. Molecular size distribution of inulin (A) and SAX (B) from high performance size exclusion chromatography with refractive index detector (HPSEC-RI). Samples were suspended into deionized water with final concentration of 1% and injected into a Sephacryl S500HR column (Amersham Biosciences, Piscataway, NJ, USA). The molecular weight (MW) was estimated from a calibration curve generated using maltodextrin standards.

Supplementary Figure 2. OTU relative abundances across all lineages. Each shade represents a different OTU and are colored according to phylum (*Proteobacteria,* blue; *Bacteroidetes*, red; *Firmicutes*, green; and *Actinobacteria*, gray). Rare OTUs (< 0.2% relative abundance in total) are combined together in Others (black).

Supplementary Figure 3. Schematic outline of 2 experiments.

## Reference

1. O’Hara AM, Shanahan F. 2006. The gut flora as a forgotten organ. EMBO Rep 7:688– 693.

2. Wong JM, De Souza R, Kendall CW, Emam A, Jenkins DJ. 2006. Colonic health: fermentation and short chain fatty acids. J Clin Gastroenterol 40:235–243.

3. Braaten JT, Wood PJ, Scott FW, Wolynetz MS, Lowe MK, Bradley-White P, Collins MW. 1994. Oat beta-glucan reduces blood cholesterol concentration in hypercholesterolemic subjects. Eur J Clin Nutr 48:465–474.

4. Giacco R, Parillo M, Rivellese AA, Lasorella G, Giacco A, D’episcopo L, Riccardi G. 2000. Long-term dietary treatment with increased amounts of fiber-rich low-glycemic index natural foods improves blood glucose control and reduces the number of hypoglycemic events in type 1 diabetic patients. Diabetes Care 23:1461–1466.

5. Cani PD, Neyrinck AM, Fava F, Knauf C, Burcelin RG, Tuohy KM, Gibson GR, Delzenne NM. 2007. Selective increases of bifidobacteria in gut microflora improve high-fat-diet-induced diabetes in mice through a mechanism associated with endotoxaemia. Diabetologia 50:2374–2383.

6. Mosca A, Leclerc M, Hugot JP. 2016. Gut microbiota diversity and human diseases: should we reintroduce key predators in our ecosystem? Front Microbiol 7:455.

7. Abrahamsson TR, Jakobsson HE, Andersson AF, Björkstén B, Engstrand L, Jenmalm MC. 2014. Low gut microbiota diversity in early infancy precedes asthma at school age. Clin Exp Allergy 44:842–850.

8. Näpflin K, Schmid-Hempel P. 2018. High Gut Microbiota Diversity Provides Lower Resistance against Infection by an Intestinal Parasite in Bumblebees. Am Nat 192:131– 141.

9. Sommer F, Anderson JM, Bharti R, Raes J, Rosenstiel P. 2017. The resilience of the intestinal microbiota influences health and disease. Nat Rev Microbiol 15:630.

10. Hamaker BR, Tuncil YE. 2014. A perspective on the complexity of dietary fiber structures and their potential effect on the gut microbiota. J Mol Biol 426:3838–3850.

11. Cantarel BL, Coutinho PM, Rancurel C, Bernard T, Lombard V, Henrissat B. 2008. The Carbohydrate-Active EnZymes database (CAZy): an expert resource for glycogenomics. Nucleic Acids Res 37:D233–D238.

12. Tuncil YE, Nakatsu CH, Kazem AE, Arioglu-Tuncil S, Reuhs B, Martens EC, Hamaker BR. 2017. Delayed utilization of some fast-fermenting soluble dietary fibers by human gut microbiota when presented in a mixture. J Funct Foods 32:347–357.

13. Gause GF. 1932. Experimental Studies on the Struggle for Existence: I. Mixed Population of Two Species of Yeast. J Exp Biol 9:389–402.

14. Hardin G. 1960. The Competitive Exclusion Principle. Science 131:1292–1297.

15. Abbott DW, van Bueren AL. 2014. Using structure to inform carbohydrate binding module function. Curr Opin Struct Biol 28:32–40.

16. McNulty NP, Wu M, Erickson AR, Pan C, Erickson BK, Martens EC, Pudlo NA, Muegge BD, Henrissat B, Hettich RL, Gordon JI. 2013. Effects of Diet on Resource Utilization by a Model Human Gut Microbiota Containing Bacteroides cellulosilyticus WH2, a Symbiont with an Extensive Glycobiome. PLOS Biol 11:e1001637.

17. Luis AS, Briggs J, Zhang X, Farnell B, Ndeh D, Labourel A, Baslé A, Cartmell A, Terrapon N, Stott K, Lowe EC, McLean R, Shearer K, Schückel J, Venditto I, Ralet M- C, Henrissat B, Martens EC, Mosimann SC, Abbott DW, Gilbert HJ. 2018. Dietary pectic glycans are degraded by coordinated enzyme pathways in human colonic Bacteroides. Nat Microbiol 3:210.

18. Lenski RE. 2017. Convergence and Divergence in a Long-Term Experiment with Bacteria. Am Nat 190:S57–S68.

19. Rycroft CE, Jones MR, Gibson GR, Rastall RA. 2001. A comparative in vitro evaluation of the fermentation properties of prebiotic oligosaccharides. J Appl Microbiol 91:878–887.

20. Walters W, Hyde ER, Berg-Lyons D, Ackermann G, Humphrey G, Parada A, Gilbert JA, Jansson JK, Caporaso JG, Fuhrman JA. 2016. Improved bacterial 16S rRNA gene (V4 and V4-5) and fungal internal transcribed spacer marker gene primers for microbial community surveys. Msystems 1:e00009–15.

21. Pokusaeva K, Fitzgerald GF, van Sinderen D. 2011. Carbohydrate metabolism in Bifidobacteria. Genes Nutr 6:285–306.

22. Van den Abbeele P, Van De Wiele T, Possemiers S. 2011. Prebiotic effect and potential health benefit of arabinoxylans. Agro Food Ind Hi Tech 22:9.

23. Gascoyne DJ, Theodorou MK. 1988. Consecutive batch culture—a novel technique for the in vitro study of mixed microbial populations from the rumen. Anim Feed Sci Technol 21:183–189.

24. Cheng YF, Edwards JE, Allison GG, Zhu W-Y, Theodorou MK. 2009. Diversity and activity of enriched ruminal cultures of anaerobic fungi and methanogens grown together on lignocellulose in consecutive batch culture. Bioresour Technol 100:4821– 4828.

25. Chung WSF, Walker AW, Vermeiren J, Sheridan PO, Bosscher D, Garcia-Campayo V, Parkhill J, Flint HJ, Duncan SH. 2019. Impact of carbohydrate substrate complexity on the diversity of the human colonic microbiota. FEMS Microbiol Ecol 95.

26. Shoaib M, Shehzad A, Omar M, Rakha A, Raza H, Sharif HR, Shakeel A, Ansari A, Niazi S. 2016. Inulin: Properties, health benefits and food applications. Carbohydr Polym 147:444–454.

27. Lu J, Li Y, Gu G, Mao Z. 2005. Effects of molecular weight and concentration of arabinoxylans on the membrane plugging. J Agric Food Chem 53:4996–5002.

28. Konopka A, Lindemann S, Fredrickson J. 2015. Dynamics in microbial communities: unraveling mechanisms to identify principles. ISME J 9:1488–1495.

29. Gibson GR, Hutkins R, Sanders ME, Prescott SL, Reimer RA, Salminen SJ, Scott K, Stanton C, Swanson KS, Cani PD, Verbeke K, Reid G. 2017. Expert consensus document: The International Scientific Association for Probiotics and Prebiotics (ISAPP) consensus statement on the definition and scope of prebiotics. Nat Rev Gastroenterol Hepatol 14:491–502.

30. Meyer D, Stasse-Wolthuis M. 2009. The bifidogenic effect of inulin and oligofructose and its consequences for gut health. Eur J Clin Nutr 63:1277.

31. Vandeputte D, Kathagen G, D’hoe K, Vieira-Silva S, Valles-Colomer M, Sabino J, Wang J, Tito RY, De Commer L, Darzi Y. 2017. Quantitative microbiome profiling links gut community variation to microbial load. Nature 551:507.

32. Bettler J, Euler AR. 2006. An evaluation of the growth of term infants fed formula supplemented with fructo-oligosaccharide. Int J Probiotics Prebiotics 1:19.

33. Calame W, Weseler AR, Viebke C, Flynn C, Siemensma AD. 2008. Gum arabic establishes prebiotic functionality in healthy human volunteers in a dose-dependent manner. Br J Nutr 100:1269–1275.

34. Scott KP, Gratz SW, Sheridan PO, Flint HJ, Duncan SH. 2013. The influence of diet on the gut microbiota. Pharmacol Res 69:52–60.

35. Sheridan PO, Martin JC, Lawley TD, Browne HP, Harris HM, Bernalier-Donadille A, Duncan SH, O’Toole PW, Scott KP, Flint HJ. 2016. Polysaccharide utilization loci and nutritional specialization in a dominant group of butyrate-producing human colonic Firmicutes. Microb Genomics 2.

36. Hoeflinger JL, Davis SR, Chow J, Miller MJ. 2015. In vitro impact of human milk oligosaccharides on Enterobacteriaceae growth. J Agric Food Chem 63:3295–3302.

37. Hidaka H, Eida T, Takizawa T, Tokunaga T, Tashiro Y. 1986. Effects of fructooligosaccharides on intestinal flora and human health. Bifidobact Microflora 5:37–50.

38. Hartemink R, Van Laere KMJ, Rombouts FM. 1997. Growth of enterobacteria on fructo-oligosaccharides. J Appl Microbiol 83:367–374.

39. Schouler C, Taki A, Chouikha I, Moulin-Schouleur M, Gilot P. 2009. A genomic island of an extraintestinal pathogenic Escherichia coli strain enables the metabolism of fructooligosaccharides, which improves intestinal colonization. J Bacteriol 191:388–393.

40. Dodd D, Mackie RI, Cann IKO. 2011. Xylan degradation, a metabolic property shared by rumen and human colonic Bacteroidetes. Mol Microbiol 79:292–304.

41. Zhang M, Chekan JR, Dodd D, Hong P-Y, Radlinski L, Revindran V, Nair SK, Mackie RI, Cann I. 2014. Xylan utilization in human gut commensal bacteria is orchestrated by unique modular organization of polysaccharide-degrading enzymes. Proc Natl Acad Sci 111:E3708–E3717.

42. Nie X, Martens E, Xiao Y, Reuhs B, Hamaker B. 2016. Substrate Structure-dependent Growth of Bacteroides Xylanisolvens XB1A on Corn Arabinoxylan Fragments in Pure and Mixed Culture Environments. FASEB J 30:683.7-683.7.

43. Rogowski A, Briggs JA, Mortimer JC, Tryfona T, Terrapon N, Lowe EC, Baslé A, Morland C, Day AM, Zheng H. 2015. Glycan complexity dictates microbial resource allocation in the large intestine. Nat Commun 6:7481.

44. Amir I, Bouvet P, Legeay C, Gophna U, Weinberger A. 2014. Eisenbergiella tayi gen. nov., sp. nov., isolated from human blood. Int J Syst Evol Microbiol 64:907–914.

45. Van den Abbeele P, Belzer C, Goossens M, Kleerebezem M, De Vos WM, Thas O, De Weirdt R, Kerckhof F-M, Van de Wiele T. 2013. Butyrate-producing Clostridium cluster XIVa species specifically colonize mucins in an in vitro gut model. ISME J 7:949.

46. Rumpagaporn P, Reuhs BL, Kaur A, Patterson JA, Keshavarzian A, Hamaker BR. 2015. Structural features of soluble cereal arabinoxylan fibers associated with a slow rate of in vitro fermentation by human fecal microbiota. Carbohydr Polym 130:191–197.

47. Pettolino FA, Walsh C, Fincher GB, Bacic A. 2012. Determining the polysaccharide composition of plant cell walls. Nat Protoc 7:1590.

48. Xu J, Chen D, Liu C, Wu X-Z, Dong C-X, Zhou J. 2016. Structural characterization and anti-tumor effects of an inulin-type fructan from Atractylodes chinensis. Int J Biol Macromol 82:765–771.

49. Chen M-H, Dien BS, Vincent ML, Below FE, Singh V. 2014. Effect of harvest maturity on carbohydrates for ethanol production from sugar enhanced temperate$\times$ tropical maize hybrid. Ind Crops Prod 60:266–272.

50. Tuncil YE, Thakkar RD, Marcia ADR, Hamaker BR, Lindemann SR. 2018. Divergent short-chain fatty acid production and succession of colonic microbiota arise in fermentation of variously-sized wheat bran fractions. Sci Rep 8:16655.

51. Ferrera I, Massana R, Balagué V, Pedrós-Alió C, Sánchez O, Mas J. 2010. Evaluation of DNA extraction methods from complex phototrophic biofilms. Biofouling 26:349–357.

52. Lindemann SR, Moran JJ, Stegen JC, Renslow RS, Hutchison JR, Cole JK, Dohnalkova AC, Tremblay J, Singh K, Malfatti SA. 2013. The epsomitic phototrophic microbial mat of Hot Lake, Washington: community structural responses to seasonal cycling. Front Microbiol 4:323.

